# Increased Activation in the Fusiform Face Area to Greebles is a Result of Expertise Training, Not by its Face-likeness

**DOI:** 10.1101/2021.09.16.460714

**Authors:** Kuo Liu, Chiu-Yueh Chen, Le-Si Wang, Chun-Chia Kung

## Abstract

In 2011, Brants, Wagemans, & Op de Beeck (JOCN 23:12, pp. 3949-3958) trained eight individuals to become Greeble experts, and found neuronal inversion effects [NIEs; i.e., higher Fusiform Face Area (FFA) activity for upright, rather than inverted Greebles]. These effects were also found for faces, both before and after training. By claiming to have replicated the seminal Greeble training study (i.e., Gauthier, Tarr, Anderson, Skudlarski, & Gore, 1999, *Nat Neurosci, 2*, 568-573), Brants et al. interpreted these results as participants viewing Greebles as faces throughout training, contrary to the original argument of subjects becoming Greeble experts only *after* training. However, such a claim presents two issues. First, the behavioral training results of Brants et al. did not replicate those of Gauthier et al (1999), raising concerns of whether the right training regime had been adopted. Second, both a literature review and meta-analysis of NIE in the FFA suggest its unreliability as an index of face(-like) processing. To empirically evaluate these issues, the present study compared two documented training paradigms (i.e., Gauthier & Tarr, 1997, *Vision Res, 37*, 1673-1682; and Gauthier, Williams, Tarr, & Tanaka, 1998, *Vision Res, 38*, 2401-2428) and explored their impact on the FFA. The results showed significant increases in the FFA for Greebles, and a clear neural “adaptation” (i.e., decreased activity for faces following Greebles, but not following non-face objects, in the FFA) both *only* in the Gauthier97 group, and *only after* training, reflecting clear modulation of expertise following “appropriate” training. In both groups, no clear NIE for faces nor Greebles were found. Collectively, these data invalidate the two assumptions behind the Brants et al. findings, and provide not only the updated support, but also the new evidence, for the perceptual expertise hypothesis of FFA.

## INTRODUCTION

Face recognition is an enduring and intriguing research topic in psychology and visual neuroscience. This is partly because of its importance to our social lives and species survival (Rivolta, 2014), but also because it showcases the intricate interplay between “nature” and “nurture” (Arcaro, Schade, Vincent, Ponce, & Livingstone, 2017; Sugita, 2008). Since the first uses of PET and fMRI in research, studies in face recognition have identified several related brain regions (Haxby et al., 1996; Sergent, Ohta, & Macdonald, 1992). However, it was only in 1997, when the term Fusiform Face Area (FFA) was coined (Kanwisher, McDermott, & Chun, 1997) that investigation of this area began in earnest. Since then, two opposing views (i.e., face specificity and perceptual expertise) surfaced; a debate that continues to this day (Gauthier, 2017; Kanwisher, 2000, 2017; Tarr & Gauthier, 2000). As claimed by supporters of the domain specific hypothesis, neural processing in the FFA is specific to faces (or face-like objects), is less recruited by other objects, and is more likely to be inherited by genetic endowment. Conversely, proponents of the perceptual expertise hypothesis assert that experience plays an important role in face processing mechanisms. Faces are something most humans see every day from birth, making it incredibly likely that life-long experience with faces underpins our species’ superb recognition capability. If this is true, the FFA should not only be activated in response to faces, but also in response to “expert” object categories, such as car experts viewing cars or radiologists viewing X-ray images (Brants, Wagemans, & Op de Beeck, 2011; Gauthier, Skudlarski, Gore, & Anderson, 2000).

To test and verify the expertise hypothesis, Gauthier and Tarr (1997) created Greebles—an artificial object set, for which they trained participants to recognize at various levels (i.e., sex, family, and individual). Participants were familiarized with various Greebles and fMRI-scanned before and after 8-10 training sessions. In one seminal fMRI study (Gauthier, Tarr, Anderson, Skudlarski, & Gore, 1999), three lines of evidence jointly supported the expertise hypothesis. Specifically, (a) decreased response times (RTs) for verification tasks with Greebles, to the point of statistical insignificance between RTs for identifying Greebles at the family level versus at the individual level; (b) comparisons of the FFA in “Faces vs. Objects” prior to training, and in both “Faces vs. Objects” and “Greebles vs. Objects” after training, suggesting that training “drove” Greeble selectivity in the FFA; and (c) comparison of activation levels before and after training found a neural inversion effect (NIE) closely associated with training (i.e., the sum of t-values in the right FFA for upright vs. inverted faces decreased, while Greebles concurrently increased, and the gap of summed t-values between faces and Greebles was significant before training, but not after training). Along with studies of natural experts (Gauthier, Skudlarski, et al., 2000; Xu, 2005), the perceptual expertise hypothesis has received decent support (Bukach, Gauthier, & Tarr, 2006). However, upon follow-up, sometimes heated exchanges have occurred (Gauthier & Bukach, 2007; Grill-Spector, Knouf, & Kanwisher, 2004; McKone & Kanwisher, 2005; McKone, Kanwisher, & Duchaine, 2007; Op de Beeck & Baker, 2010a, 2010b).

Among objections to the expertise hypothesis of the FFA, the most relevant was a study by Brants et al. (2011), who claimed that the FFA’s increased response to Greebles was due to their resemblance to faces. Brants et al. claimed that, by replicating Gauthier’s classic training paradigm (Gauthier & Tarr, 1997), they found NIEs in the FFA, as assessed by significantly higher blood-oxygen-level dependent (BOLD) activity for upright than for inverted stimuli (either faces or Greebles), both before and after training. In addition, Brants et al. found that either encouraging or discouraging subjects from seeing Greebles as “living individuals” or “objects” did not matter; all subjects reported perceiving Greebles as “face-like” after training. Based on the post-training interview and the NIE observed for faces and Greebles, both before and after training, Brants et al. concluded that it was Greebles’ perceived face-likeness, rather than acquired expertise, that drove the NIE, which presumably reflected face-selectivity for the FFA.

Upon closer inspection, however, four inconsistencies (two behavioral, and two fMRI) between Brants et al. (2011) and Gauthier et al. (1999) could be identified. First, as the basic premise of a successful replication is an almost identical, or at least comparable, behavioral result (typically a prerequisite for the later fMRI findings), Brants et al. (2011) reported mean RTs for accurate verification trials of ~1000 ms. This was nearly double the RTs (~500 ms) reported in the original study (Figure 3 of Gauthier & Tarr, 1997, p. 1677; Gauthier et al., 1999). Although Brants et al. (2011, p. 3951) stated that “All aspects of the procedure were modeled after Gauthier et al. (1997), except the manipulation of inducing face-likeness for half of the participants,” they nevertheless also stated “ten Greebles remained unknown throughout the training” (p. 3951), which happened in neither the Gauthier and Tarr (1997), nor the Gauthier et al. (1999) studies. In fact, such a “half-trained, half-untrained” arrangement of Greebles more resembles that found in Gauthier, Williams, Tarr, and Tanaka (1998), where the verification RTs for these two categories differed significantly (ibid, Figure 4B, pp. 2407), and the mean verification RT of ~1050 ms (averaging the former mean of ~800ms and the later mean of ~1300ms) was also more consistent with those reported by Brants et al. (2011, Figure 3B, p. 3953). These observations and reasoning clearly suggest that Brants et al. (2011) adopted a different, or variant of the training paradigm (i.e., Gauthier et al. 1998), than those originally used in Gauthier and Tarr (1997) and Gauthier et al. (1999). However, it would still be necessary to empirically compare the effects of different expertise training in the FFA/inferior occipitotemporal cortex to draw a final conclusion. Second, Brants et al. (2011) used the “statistical equivalence between mean verification RTs between family and individual level” (Gauthier & Tarr, 2002) as the sole criterion for subjects to reach perceptual expertise. However, according to Tanaka and Gauthier (1997), two other implicit criteria also need to be met to jointly define “perceptual expertise”: (a) that the RTs between Greeble verifications at the family vs. those at the individual levels are supposed to be large and/or significantly different, with the latter usually longer than those of the former; and (b) the RT differences would monotonically decrease to the point of near convergence, or statistical insignificance. With only one criterion (i.e., statistical insignificance, as depicted by three arrows; session 1, 2, and 8, respectively; Brants et al., 2011, Figure 3B, p. 3953), it is unimaginable that experts and novices could sometimes be separated, and sometimes not.

Third, while both Gauthier et al. (1999) and Brants et al. (2011) reported behavioral training results and comparisons of inversion effects in the FFA for faces and Greebles, the former provided additional contrast images of category-selective regions. As shown in Gauthier et al. (1999, Figure 4, p. 571), before training, the FFA could only be identified in the “Face vs. Object” contrast. After training however, contrasts of both “Face vs. Object” and “Greeble vs. Object” revealed the same FFA. If, according to Brants et al. (2011), participants interpreted Greebles as faces throughout training, an further convincing support of such claim would be to similary show FFA activations for “Face/Greeble vs. Object” contrast both before *and* after training. While this was not seen in Brants et al. (2011), a counterargument to the “Greebles look like faces” claim could be that: since most of us do sometimes consider Greebles “face-like” *prior* to training, the fact that “Greeble vs. Object” contrast revealed FFA only *after*, but not *before*, training, as revealed in Gauthier et al. (1999), strongly suggest the necessity of sufficient training to shift processing from the basic to subordinate level (Gauthier, Anderson, Tarr, Skudlarski, & Gore, 1997; Gauthier, Tarr, et al., 2000) as the crucial factor driving increased FFA activity for Greebles, not by Greebles’ face-resemblance alone. Lastly, as both Gauthier et al. (1999) and Brants et al. (2011) observed NIEs in the FFA as the indispensable evidence for their respective claim, the reliability of NIE@FFA as an index of face processing was hardly examined. Furthermore, the above two studies adopted different dependent measures: summed t-values (Gauthier et al., 1999)over and average percent signal change, or PSC, across (Brants et al., 2011), all FFA voxels. As behavioral inversion effects may not be face-specific (Rossion & Gauthier, 2002; Valentine, 1988), our extant literature search by keywords like “FFA” and “face inversion,” rendered 14 published articles (shown in Table 1). Not only did these articles show surprisingly inconsistent NIE effects at FFA, further funnel plot and meta-analyses also did not yield significant differences, suggesting that the NIE@FFA was never unanimously one-sided (larger for upright orientation). In light of the obvious unreliability of NIEs as an indicator of face-specific processing, it may be worthwhile to look for an alternative neuronal measure of face/expertise processing.

**Table 1:** A summary literature of the neural inversion effect of face in FFA. 18 articles, 7 of them found a positively (i.e. larger activity for upright than for inverted faces) significant NIE (Green cells and points), 9 of them did not find significant NIE (Yellow cells points, with 5 of them in darker yellow colors, indicating their lack of statistical, t/p/F, values) and 2 found a reverse NIE (Blue cells points).

| Study | fMRI design | Number of subjects | (NIE) for faces |
| --- | --- | --- | --- |
| Kanwisher et al. (1998) | blocked | 10 | $F(1,9) = 10.6, *p < 0.01$ |
| Yovel et al. (2004) | blocked | 17 | $F(1,13) = 16.5, *p < .001$ |
| Yovel et al. (2005) | blocked+er_fMRI | 21+14 | $t(30) = 4.67, *p < .001$ |
| Passarotti et al. (2007) | blocked | 3 groups, N=30 | Pos. NIE for adults ($F(1,27) = 4.81, *p < .04$) |
| Rhodes et al. (2009) | blocked | 16 | $F(3,39) = 17.34, *p < .0001, \eta^2 = .57$ |
| Brants et al. (2011) | blocked | 8 | $F(1, 7) = 12.083, *p < .05.$ |
| James et al. (2013) | blocked | 12 | $t(11) = 4.92, q < .05$ (Fig. 5) |
| Haxby et al. (1999) | blocked | 6 | Not sig. |
| Aguirre et al (1999) | slow event-related | 8 | $t(7) = .88$, not sig. |
| Epstein et al (2006) | blocked | 12 | $p > 0.15$ |
| Kung et al. (2007) | blocked | 10 | not sig. ($t(18) = 0.662, p = 0.26$) |
| Strother et al. (2011) | blocked | 12 | not sig. (judged by Fig. 4 & 5a) |
| Bilalic et al (2011) | blocked | 15 (7 experts and 8 novices. in Exp1) | not sig. (stats unreported. Fig. 1D) |
| Grotheer et al. (2014) | blocked | 26 | $F(1,25) = 0.09, (*p < .05)$ |
| Matsuyoshi et al. (2015) | event-related | 32 | Not sig. (Fig. 4). |
| Chou et al. (2021) | event-related | 32 | Not sig (Fig. 3B). |
| Bookheimer et al. (2008) | blocked | 12 (ASD) and 12 ND | Reversed NIE $F(1,154) = 38.73, p < .001$ |
| Passarotti et al. (2007) | blocked | 3 groups, N=30 | Reversed NIE for children ($F(1,10) = 11.86, p < .002$) |

With these four comments, the purpose of the present study was to revisit the two Greeble training regimes: (Gauthier & Tarr, 1997; Gauthier et al., 1999) and (Brants et al., 2011; Gauthier et al., 1998), and to compare their differential interactions with the FFA. To show that FFA selectivity is not driven by Greebles’ face-likeness (but more likely by subordinate-level processing), we chose asymmetric Greebles to minimize the effect of symmetry. If all putative Greeble selectivity in the FFA could still be observed for the asymmetric version, stimulus symmetry (i.e., a potential factor behind face likeness) becomes less important in shaping the selectivity of the FFA. Additionally, in the current experiment both the FFA and lateral object complex (LOC) would be independently localized (c.f., Brants et al., 2011). Other experimental procedures, including the Greebles-Objects-Faces (GFO) tasks and the face/Greeble inversions manipulation, were kept as closely matched with those in both Gauthier et al. (1999) and Brants et al. (2011).

The domain specific hypothesis predicts that Greeble training does not exert any effect on FFA, but could be on another object selective area (e.g., LOC). In contrast, the perceptual expertise hypothesis predicts that “Greeble vs. Object” contrast should reveal the FFA *only after* training. We predicted that different training regimes would also yield different response profiles, and different Greeble selectivities in the FFA accordingly. Regarding NIEs at the FFA, we are equivocal about whether replication of previous findings were easy, since both the meta-analyses and forest plots suggest mixed NIE results. Lastly, the expertise hypothesis predicts that if the FFA becomes similarly responsive to faces and Greebles after training, we would expect neural adaptation (Grill-Spector, Kushnir, Edelman, Itzchak, & Malach, 1998; Grill-Spector & Malach, 2001) or competition in the FFA: for example, reduced FFA activities for faces right after Greebles, but only after training, and only under the right training regime.

## METHODS

### Behavioral Experiment

#### Participants

Sixteen NCKU undergraduate and graduate students (nine females), all with normal or corrected-to-normal vision, were recruited. After signing the informed consent approved by the NCKU Research Ethics Committee (REC case number 103-266), participants were randomly assigned to two separate training paradigms (*n* = 8 in each), which included three fMRI scans (before, during/halfway, and after training), and 10 sessions of behavioral training in between, lasting approximately two weeks. At the end of training, all participants received a cash payment of ~NT$ 6000.

#### Materials

The 60 Greeble exemplars (Figure 1), are categorized by two genders, five families, and six individual levels (also see Gauthier & Tarr, 1997, Figure 1, p. 1674). Gender is defined by Greebles’ upward or downward “appendages” (i.e., protrusions on the top and main body), the family by the five different body shapes, and the individual by the unique combination of body shape and the appendages. Also shown in Figure 1 are meaningless syllables assigned to these asymmetric Greebles as their respective gender, family, and individual name, with all the initial letters different for the convenience of responses in the later naming task.

**Figure 1.**
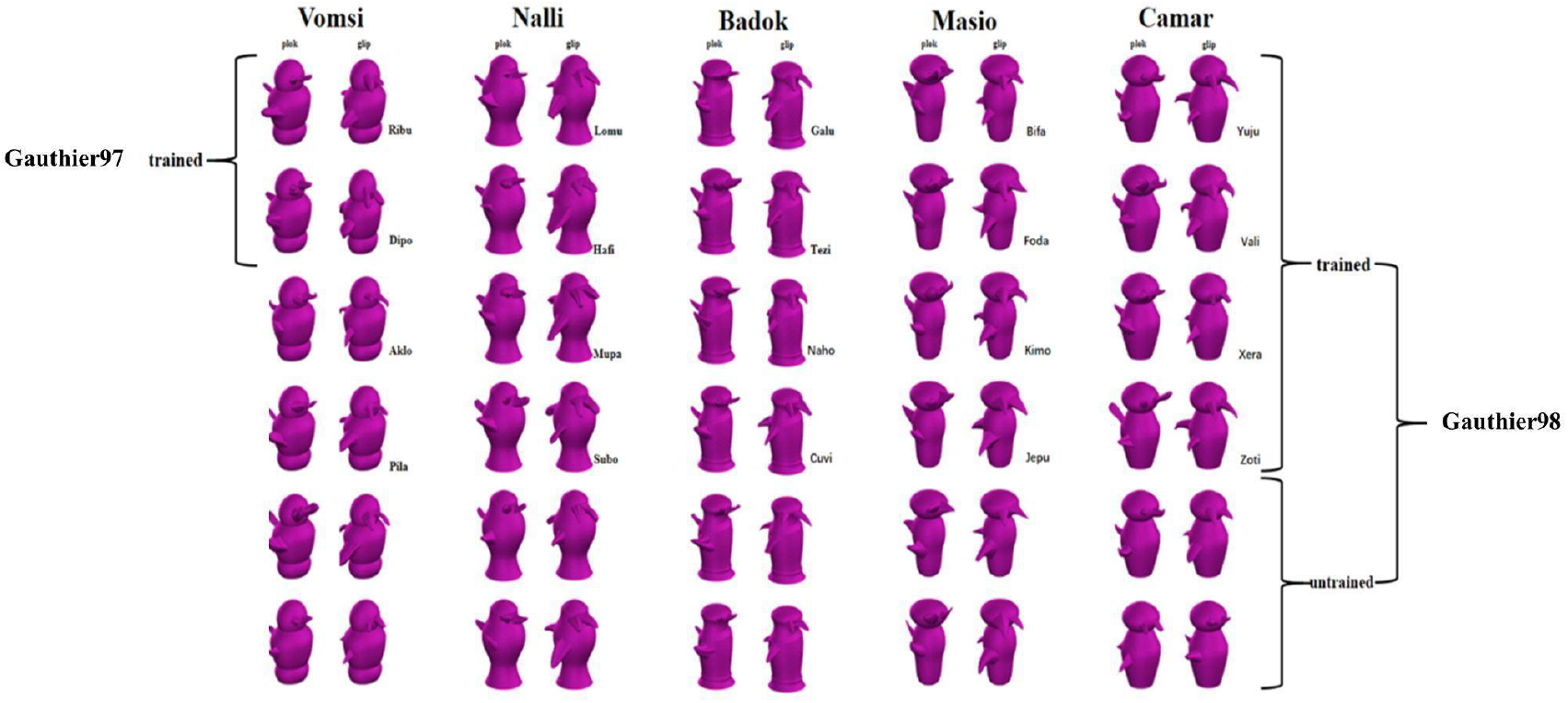
Thirty asymmetric Greebles used in the present study, with each column representing a family (a total of 5, with family names above), each family contain six individuals each (names in between the two trained views, which presented equally throughout the training process. There are three differences between Gauthier97 and Gauthier98: a. Gauthier97 have family level to learn whereas Gauthier did not; b. Gauthier97 had ten individual names to learn, while Gauthier98 had twenty. c. Gauthier98 added unnamed label in naming and verification task. All Greeble stimuli were downloaded form: http://wiki.cnbc.cmu.edu/Novel_Objects.

Both training regimes were performed in MATLAB 2010a on an iMac with the resolution of 1024*768. All Greeble models were presented in purple shade (Figure 1). The size of presented images was approximately 6.5 cm high and 3.25 cm wide (nearly identical to Brants et al., 2011). Every participant saw Greebles in the center of the screen, which was placed at a distance of 60 cm, subtending an approximate 6.2° × 3.1° visual angle.

#### Procedure

The sixteen participants were randomly divided into two groups (i.e., training regimes). The first group followed the training procedure of Gauthier et al. (1997, 1999; shortened to Gauthier97 hereafter), and the other followed those by Gauthier et al. (1998) and Brants et al. (2011, shortened to Gauthier98 hereafter). As stated in Gauthier et al. (1998, p. 2403), there were three explicit procedural differences between the Gauthier97 and Gauthier98 groups. First, Gauthier97 participants were trained to classify Greebles at all three levels (gender, family, and individual), whereas Gauthier98 only included gender- and individual-level training. Second, Gauthier98 trained participants required discrimination of 10 different Greebles (two from each gender, and five from each family), as opposed to 20 in the Gauthier98 protocol. Finally, Gauthier97 had participants perform only the verification task during training, while Gauthier98 required participants to alternate between the verification and naming task during most of the training sessions. Another outstanding, but less clearly stated, difference was the treatment of unnamed Greebles during training (also see Figure 1 of the present study). Of the 60 Greebles (or 30, if picking only one gender per family), only the chosen ten were trained at the individual level in the Gauthier97 protocol, leaving the remaining 20 Greebles at the gender and family level. In contrast, in the Gauthier98 protocol, other than the 20 trained Greebles, the remaining 10 unnamed ones were always present and mixed with 20 trained ones during training at both gender and individual levels, creating additional responses in both naming (pressing letter “u” for “unnamed”) and verification (pressing spacebar for “null” responses) tasks. We believe (explained later in Figure 9) that these are the primary reasons driving the RT differences in the verification task between Gauthier98 and Gauthier97. Except for these differences, all other aspects of the training regimes, including the trial structure (e.g., time intervals, stimuli addition order, a “beep warning sound after incorrect trials) were mimicked as closely as described in both Gauthiers’ protocols. All 16 participants (8 in each training regime) completed ten training sessions (spanning approximately 2 weeks), with each session lasting approximately 40 minutes. At the end of each naming/verification task block, mean accuracy and mean RT of accurate trials were shown on the screen. Lastly, at the beginning and end of training, we also asked each participant questions like “What do you think the Greebles look like?” and “Were the Greebles ever face-like to you at any time during training?”

For data analysis, both accuracy and RTs for correct responses across each training group of participants (*n* = 8) were averaged. Throughout the study, we used the same definition of expertise criterion (Gauthier & Tarr, 1997; Gauthier et al., 1999), specifically statistical insignificance between correct RTs at individual and family (Gauthier98)/gender (Gauthier97) level, during the verification task.

### fMRI

Several fMRI tasks were administered at various time points. Specifically, before training (1^st^), during training (2^nd^), and typically no less than 24 hours after reaching expertise criterion (3^rd^).

#### A. Greeble-Face-Object (GFO) task

Present in both Gauthier et al. (1999) and Brants et al. (2011), the GFO task is necessary for both examination of training effects and for cross-study comparisons. It requires subjects to passively view two runs (322 s; 161 volumes each) of 12 image blocks (each category repeated four times, 20 images per block, stimulus-onset asynchrony of 1 s), sandwiched between 6 s fixation periods (first fixation lasting 10 s for dummy scan), in both before- and after-training fMRIs. Unique in the current protocol, in order to assess the relative interference or neural adaptation (e.g., face stimuli presented after the Greeble block), each category was with equal probability before and after the other two categories, so that the four face blocks would be twice after Greebles and twice after objects, with the same being true for both Greebles and objects).

#### B. NIE task

Also adopted from both Gauthier et al. (1999) and Brants et al. (2011. See Figure 2, p. 3952 for illustration), each NIE run contained four blocks of sequential matching trials (eight per block), where each trial consisted of fixation (800 ms), followed by consecutive presentations of stimulus (1 s for each image, separated by a 200 s mask in between). Four category (Greebles/faces) by orientation (upright/inverted) combinations were block-randomized in each run, and fMRI data from five runs were collected both before and after the training session. Behavioral reaction time and accuracy inside the scanner were also recorded.

**Figure 2.**
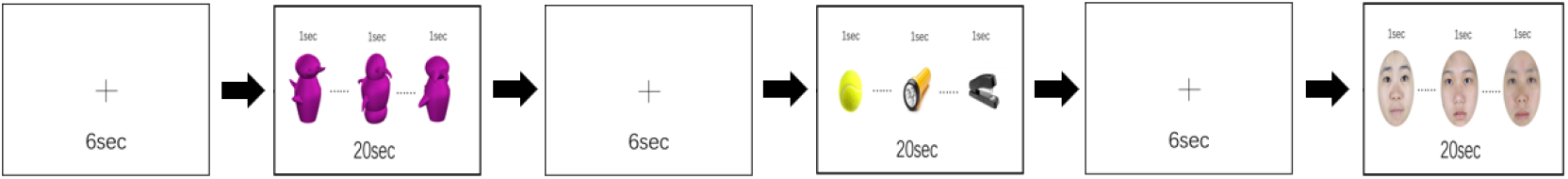
An example of GFO (Greeble, Face, Greeble) task used in fMRI scan. Stimuli were randomly presented for 1sec each during a 20sec block, while fixation blocks (6sec) in between each stimulus block. Gauthier used this tor localizer but here different localizer task Was used.

#### C. FFA Localizer task

To independently assess the differential effects of training on GFO activities, we additionally ran an independent FFA localizer (from Peelen & Downing, 2005) for each participant. Participants performed a one-back identity judgment for faces, bodies, scrambled objects, and Greebles (arranged in +ABCD+, a pseudo-random fashion) for two runs. In addition to FFA, an object-selective region (i.e., the LOC) was also circumscribed for companion analyses.

#### Scan parameters and analysis

The fMRI data were acquired from a GE750 3T scanner housed in the NCKU MRI Center, with a 32-channel head coil. Parameters were set to TR = 2000 ms, TE = 33 ms, 64 x 64 matrix (so that FOV = 19.2 x 19.2 cm), 3 mm slice thickness (without gaps, making the voxel size = 3 x 3 x 3 mm^3^), and forty axial slices, covering the whole brain.

Data was analyzed by BrainVoyager QX (BVQX v2.6) and NeuroElf 1.0 (http://neuroelf.net) under MatLab 2012a. Functional data preprocessing included slice-time correction and motion correction. The resulting functional (T2*) images were coregistered (with the normalized T1 image in the Talairach space), then transformed into 3D volume time course (VTC) data, which was then entered into a generalized linear model (GLM) and contrast analysis. Cross-session alignments of the anatomical images were performed for the first session (before training), so that all subsequent volumes could be cross-compared. For the ROI analysis, the two-run localizer data first entered the GLM and then “Faces vs. Objects” contrast applied, with the FFA defined individually and exclusively in the right hemisphere (FFA was located in the Talairach space of X: 40–50; Y: −45– −55; Z: −10– −20, with a corrected threshold of *p* < 0.5. The summed t-values and FFA averaged PSC were extracted and compared with the earlier findings.

## RESULTS

### Behavioral Results

Figure 3 summarizes the mean accuracy (left panel) and RTs (right panel) by session results of the verification tasks in the two Greebles training regimes (Gauthier97 on top). For both Gauthier97 and Gauthier98, the gender accuracy were all around 90% since the first session ([F_(1,14)_ = 0.145, p = 0.71, η^2^ = 0.01). As for the accuracy at the family and the individual levels, participants all improved with training (comparing two groups’ individual-level accuracy between the first and the last(tenth) session, F_(1,30)_ = 108.35, p <0.01, η^2^ = 0.78]). These accuracy results (Fig. 3a and 3c) suggest that all participants were able to accurately identify Greebles on the designated (gender, family, or individual) level at the end of training.

**Figure 3.**
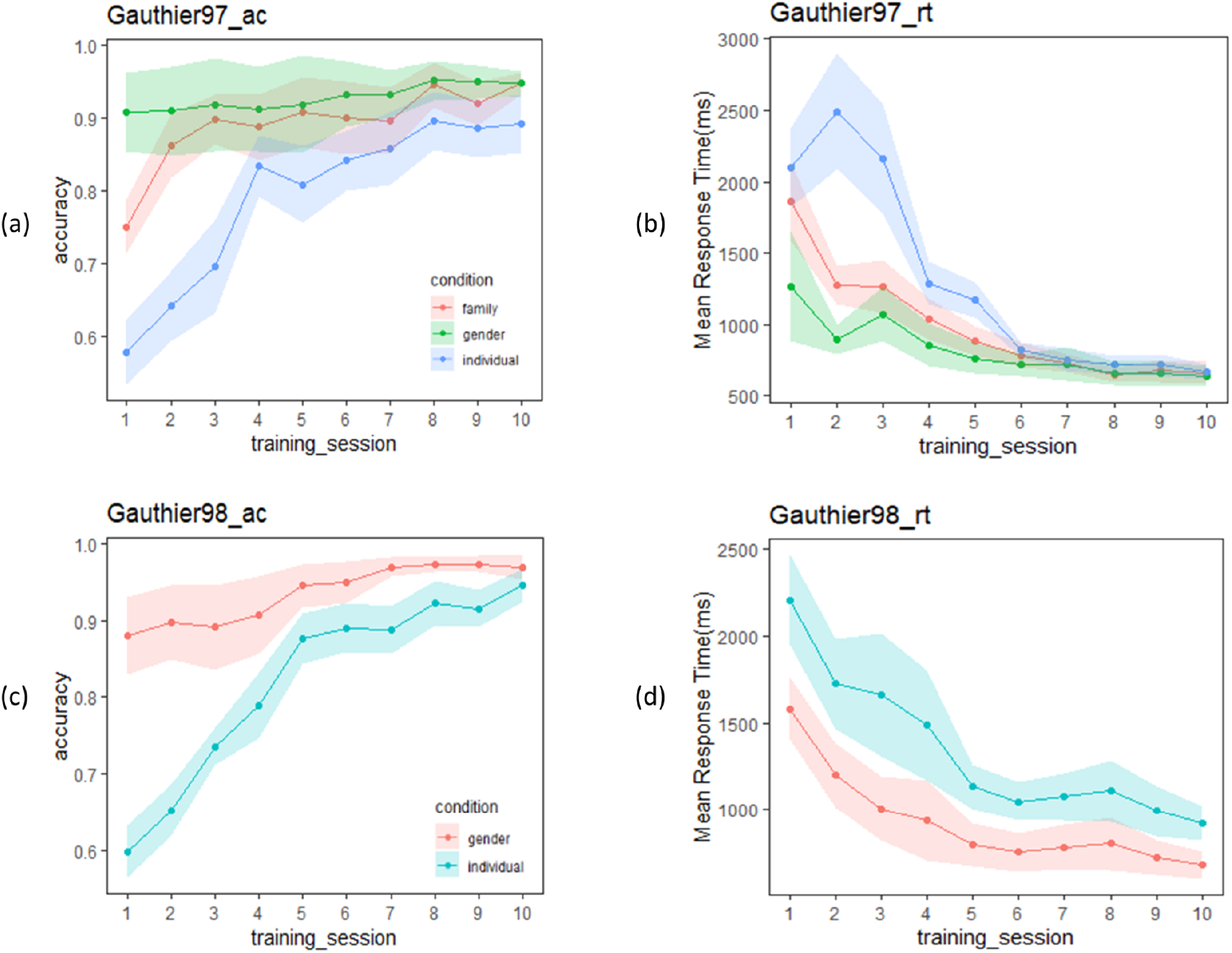
(a) Mean accuracy of Gauthier97 group (n= 8) for the three types of category level (gender, family, and individual levels) of the verification task; (b) Mean response time (in millisecond) of Gauthier97 on the same verification task. The response time of three category levels was statistically insignificant from Day 4. (c) Accuracy of Gautheir98 group (n= 7) for the two types of category level (gender and individual levels). (d) Mean response time (in millisecond) of Gauthier98 in the verification task. Note that the individual level RT combined both named and unnamed Greebles, and the similarity between our (d) with Fig. 38 (pp. 3953) of Brants, Wagemans, & Op de Beeck (2011).

In terms of RT, although in Figure 3b (Gauthier97) the mean response times in the individual level were longer than those in the family and the gender levels [the 1st ~5th session: F(2,21)= 0.07, p = 0.02~, η2 = 0.0069], with training the RT difference became closer, and finally converged from the sixth, and all the way to the 10^th^, session [the last 5 F(2,21)= 0.07, p = 0.929~0.932, η2 = .0066~0.0069]. According to the canonical definition of Greeble expertise (Gauthier & Tarr, 1997; Tanaka & Gauthier, 1997; usually focusing on the statistical insignificance between verification RTs of family and those of individual levels), our Gauthier97 subjects have become experts since the 6th session, and kept practicing for 4 more sessions.

In contrast with Fig. 3b, the Gauthier98 RT results (Fig. 3d) were quite different: not just that the mean RT were about 1000ms (compared to the mean final RT in Fig. 3b, around 500ms), there also seemed no trend of convergence throughout Gauthier98 training. Strikingly, our Fig. 3d results were very similar to the training results reported by Brants et al. (2001., Fig. 3B, pp. 3953). Inspecting their reported procedure: “*Participants were trained following the procedure used by Gauthier et al. (1997)*.” (ibid, pp. 3951), we found the match between Brants et al.’s Fig. 3B (pp. 3953) and the Fig. 3d (plus the mismatch with the Fig. 3b) of the current study lends support to the idea that Brants et al. (2001) misquoted the Gauthier98 as Gauthier97 training paradigm. Their following methodological descriptions “*The participants learned the family labels and individual names for five Greebles in the first session and then learned five more Greebles in each of the following three sessions. Ten Greebles remained unknown throughout the train, which made the tasks more difficult*,” (ibid., 3951) further verified our suspicion (of their misquote).

Two follow-up observations remain. First, since our Gauthier98 results mimicked those of Brants et al. (2001), how about the replications between our Gauthier98 results and those from the original Gauthier et al. (1998)., especially, the gender and named/unnamed individual-level training RTs? For this comparison (Fig. 4b, ibid, pp. 2407; and the similar plot of the current study, shown in the lower right corner of Fig. 5), one sees a slightly different picture: while the individual-named Greebles RTs gradually converged with the gender RTs in both studies (similar to what was found in Gauthier et al. 1998), our individual-unnamed Greeble RT, while also not converged with the mean gender RT, was gradually decreased with training (unlike what was shown in Fig. 4b, of Gauthier et al. 1998, where the unnamed RT was constantly high). Further comparison between our individual training RT plots (Fig. 5), alongside the similar Figure 5 of Gauthier et al. (1998; pp. 2408), revealed that one likely explanation was the individual variability inherent in the small samples of both the original (N=12) and the current replicated Gauthier98 (N=8) study, respectively. Overall, both studies shared the converging finding between the gender and the individual-named RTs, but no-converge between the gender and individual-unnamed RTs (Fig. 4b of Gauthier et al., 1998, and lower right of Fig. 5 in this study). This inability to converge between the two (here individual-combined and gender) levels, a signature of perceptual expertise (Tanaka and Gauthier, 1997), renders Gauthier98 training regime unfit for the behavioral training of Greeble experts.

**Figure 4.**
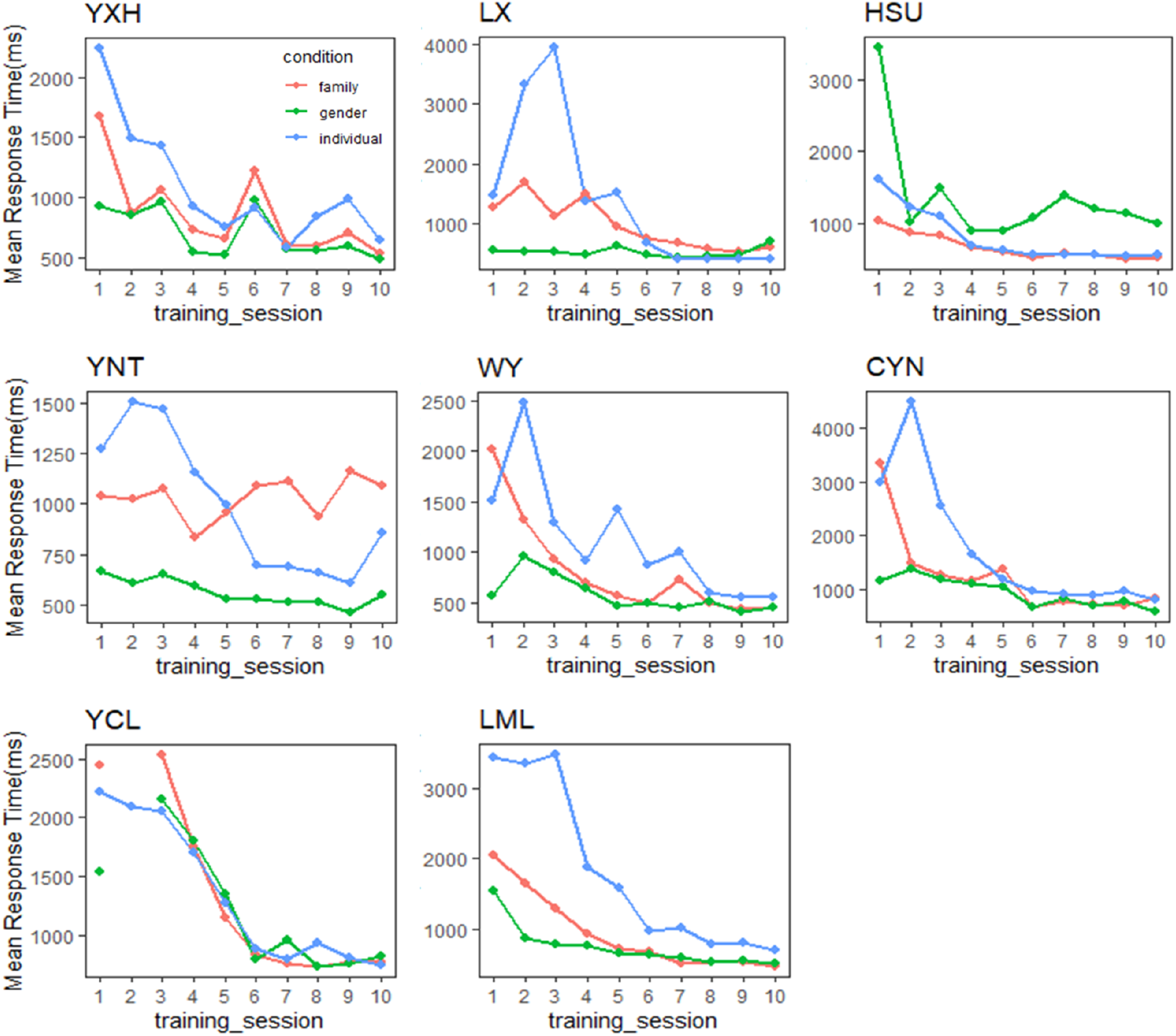
Performance in the verification task throughout training for each expert participant in the Gauthier97 group. Convergence happened among the trends of the mean response time of gender, family, and individual name. Only in the Gauthier97 group, and only after training, which reflects modulation of the expertise category following “appropriate” training regime.

**Figure 5.**
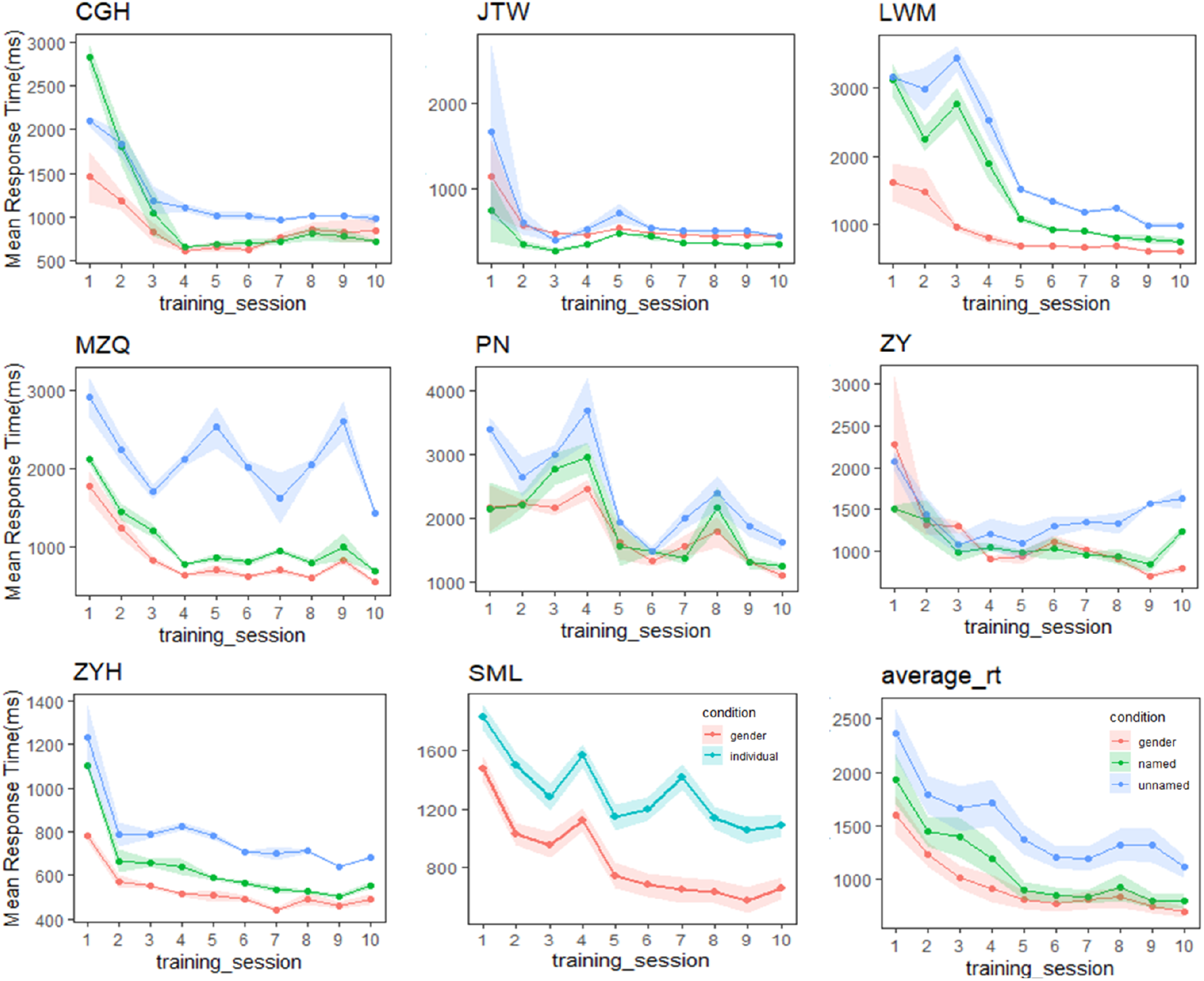
Performance in the verification task throughout training for each expert participant in the Gauthier98 group. While the individual-named Greebles RTs gradually converged with the gender RTs, our individual-unnamed Greebles also gradually decreased, but not converged, with training. One potential explanation was the individual variability inherent in the small samples. Overall, this inability to converge between the two levels renders Gauthier98 training regime unfit for the behavioral training of Greeble experts.

Second, there was an expertise-reaching RT difference between the two training regimes: in Gauthier97 it was around 500 ms at the end of expertise training; whereas in Gauthier98 the end RT hovered around 1000 ms. Such a two-fold RT difference (500 in Gauthier97 vs. 1000 ms in Gauthier98) likely resulted from the additional processing steps in Gauthier98. Fig. 9 (in discussion) lists two of the possible stages of such putative decisions Greeble trainees faced during the training processes: one upon the label presentation, and another upon Greeble presentation. With the additive factors logic (Sternberg, Markby, Wender, & Mohilner, 1969), the 500 ms gap between the two training regimes could be reasonably explained.

In sum, the converging RT results with Gauthier97 training, as shown in Figure 4, in contrast to the no-converging results with Gauthier97 paradigm, suggest that only the Gauthier97 group, and only after training, were the participants successfully achieved the criterion of perceptual expertise. We therefore concluded that the expertise training paradigm of Gauthier97 was more robust, and therefore more “appropriate”, training regime (than that of Gauthier98, or Brants et al. 2011) in generating Greeble experts.

### fMRI Results: ROI (FFA) analyses

To distinguish different accounts of the FFA activities, namely the perceptual expertise vs. the face-likeness, three proposed ROI (FFA) analyses were tested here, all between Gauthier97 and Gauthier98 training regime: (a). the comparisons among upright faces, objects, and Greebles, between before- and after-training; (b). the neuronal inversion effect (NIE): the upright vs. inverted Greebles and faces, across before-, during-, and after-training; and ©. the neural adaptation effects for both faces and Greebles, again before-vs. after-training. The perceptual expertise (PE) hypothesis of the FFA predicts that (a) FFA responses to Greebles would increase significantly from the before-to after-training, but no such effect for faces or objects, and only be so in the appropriate (e.g., Gauthier97) training paradigm; and (c) a significant increase in Faces after Greebles (e.g., neural adaptation) only after training, and under the appropriate, or Gauthier97, paradigm, as well. In contrast, FFA face specificity hypothesis, according to Brants et al. (2011), would predict: (b) a NIE for faces both before- and after-training, but not for asymmetric Greebles (given their much less, if not zero, face-likeness), given the premise that NIE being a consistent and reliable index of face-related processing, an assumption that will be examined in the meta-analysis of extant literature.

### (a) ROI (FFA) analysis

The results of FFA activities for faces, objects, and Greebles across before- and after-training, as well as for both Gauthier97 and Gauthier98, were shown in Figure 7. For the ‘Faces vs. Objects’ comparison, the main effects were found for both before- [*F*_(1,110)_ = 301.29, *p* < 0.01] and after-training [F(1,110) = 195.64, p < 0.01]. No training effect (after-vs. before-training) was found for faces in the FFA [*F*_(1,110)_ = 0.1353,*p* = .71]. Likewise, the comparison of “Greebles vs. Objects” yielded a main effect both before- [*F*_(1,110)_ = 6.17, *p* = 0.014] and after-training [*F*_(1,110)_ = 22.97, *p* < 0.01]. More importantly, in the Gauthier97 paradigm there was a significant difference for Greeble activations before- vs. after-training [*F*_(1,110)_ = 7.31, *p* < 0.01], indicating the Greeble training effect onto the FFA (see Fig. 7 left).

**Figure 7.**
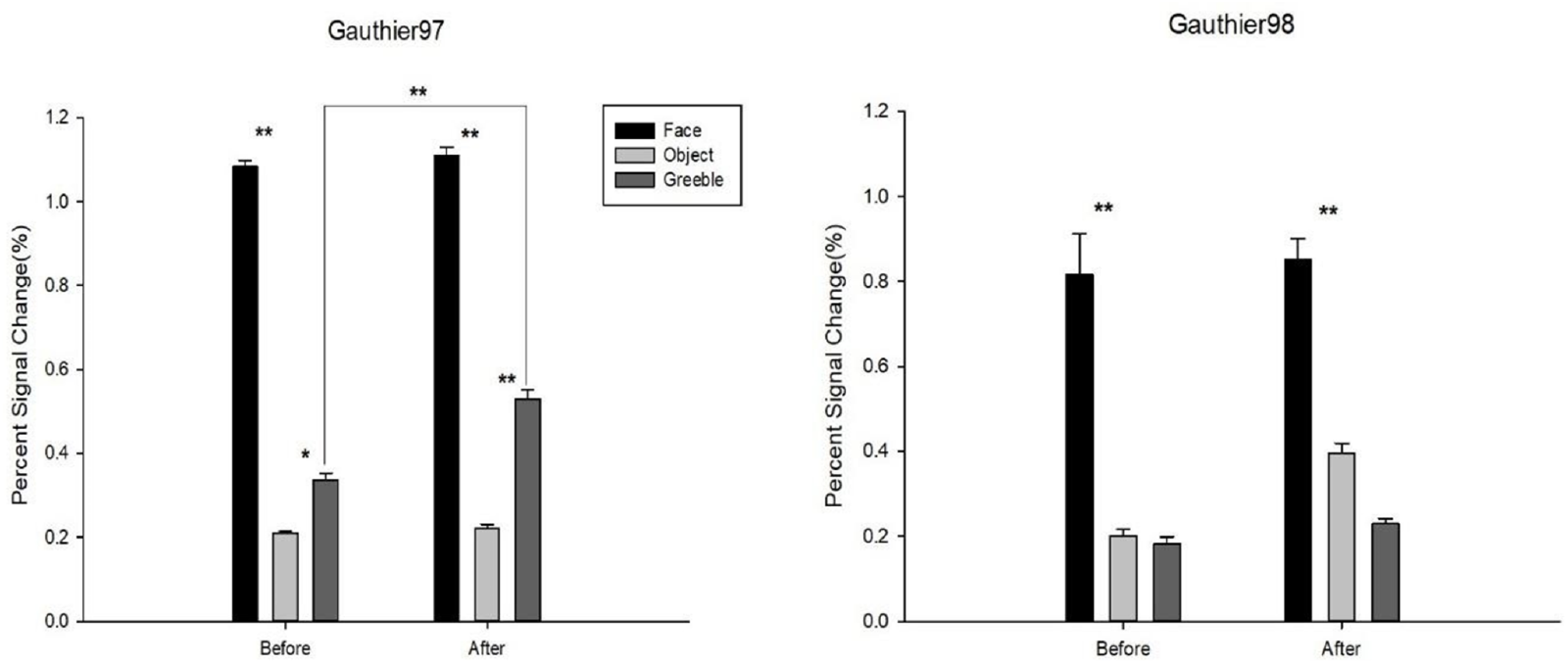
Percent signal changes for faces, objects, and Greebles in the FFA. Error bars were standard errors. Shown in left by Gauthier97 training results, right by Gauthier98. N=8 in each training group. As shown, there was a significant training effect for Greebles in the Gauthier97, but not in the Gauthier98, paradigm.

In contrast, as shown in Fig. 7 right, other than the similarly significant ‘Faces vs. Objects’ main effects both before- [*F*_(1,110)_ = 26.06, *p* < 0.01] and after-training [F_(1,110)_ = 21.12, *p* < 0.01], Gauthier98 results showed none “Greebles versus Objects” main effects across training (both *p* >.05). Because the current training regimes strictly adhered to those in both Gauthier et al. (1999) and Brants et al. (2011), with the only exception the choice of asymmetric Greebles, these results were more in line with the perceptual expertise account.

### (b) Neural Inversion Effect (NIE) in the FFA

Behavioral performance of all participants in the sequential matching task during the three fMRI sessions was shown in Fig. 8. Both groups demonstrated improvements after five sessions of training [before- vs. after-training: Gauthier97: F_(1,78)_ = 7.1324, *p* < 0.01; Gauthier98: F_(1,78)_ = 16.68405, *p* < 0.01), different from the results reported by Brants et al. (2011).

**Figure 8.**
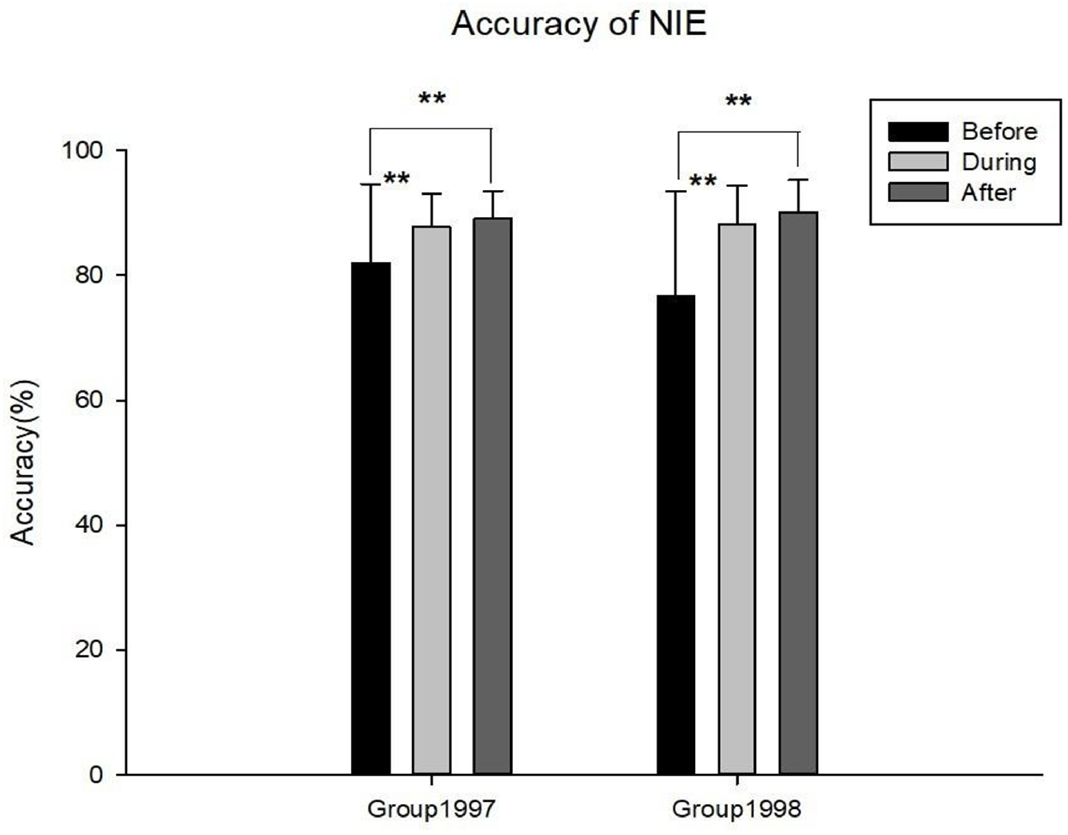
The mean correct rates of identity-matching task, either upright or inverted faces or Greebles, across three fMRI scan sessions. Error bars represent standard errors.

Mean Percent Signal Change (PSC) difference between the upright and the inverted faces or Greebles were defined as the neural inversion effect, or NIE. For faces, no significant NIEs were observed in either Gauthier97 and 98 (Figure 9a and 9c), across all three scan sessions (befor-, during-, and after-training). Out of the six possible face FIEs (three for each training group), only one (the during-training of Gauthier97) was significant [*F*_(1,126)_ = 7.83, *p* = 0.0059]. As for Greebles, none of the six NIEs was significant (see Figure 9b and 9d). These results are in sharp contrast to Brants et al. (2011), who reported significant FIEs for faces and Greebles, both before- and after-training (4 out of 4). In other words, while the present study only found one significant NIE out of 12 possible tests (8.3%), Brants et al. got 4 out of 4 (or 100%) significant NIEs!

**Figure 9.**
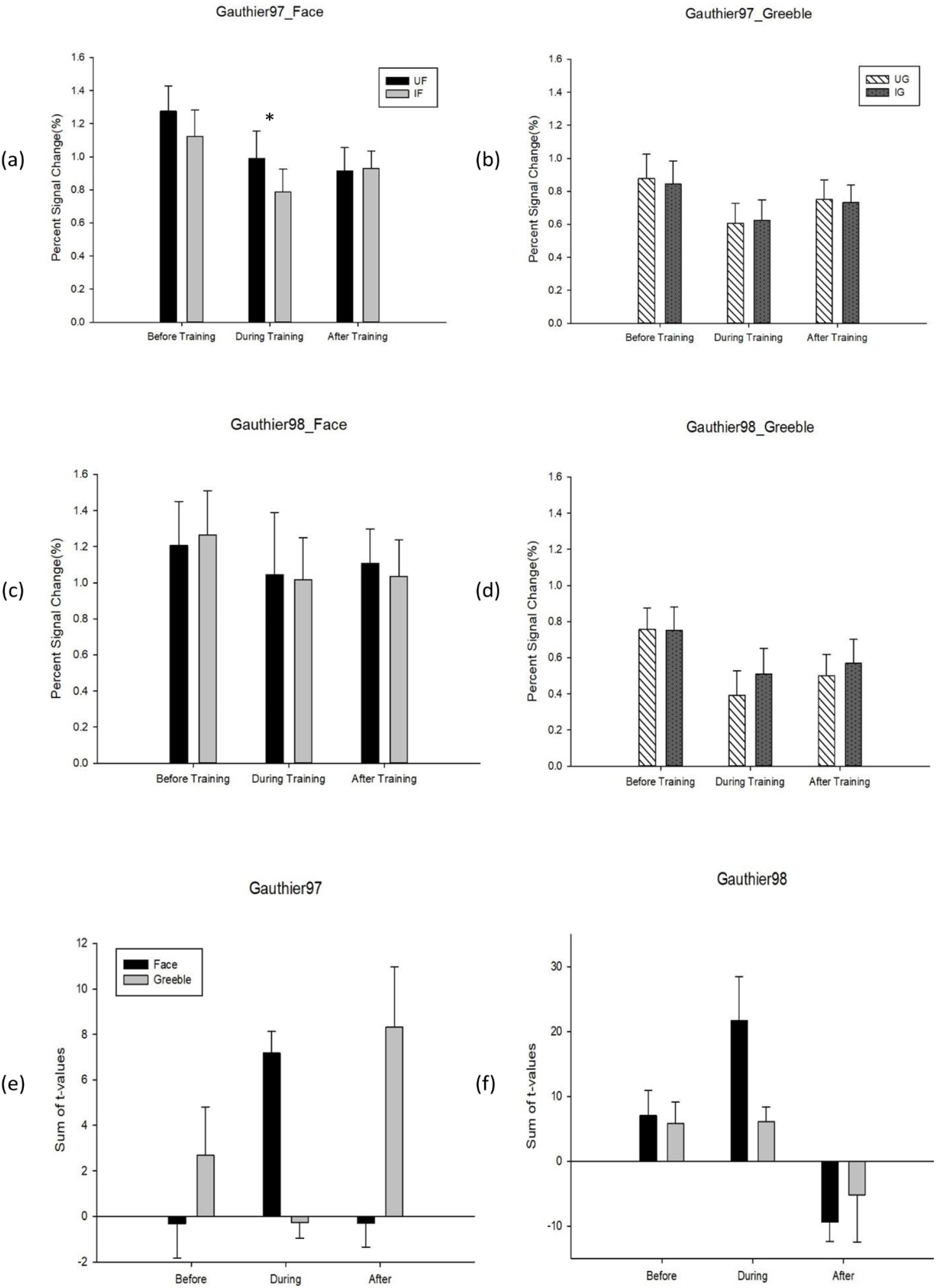
(a) FFA activities (percent signal change, or PSC) to the upright versus inverted faces; (b) the same FFA neural inversion effect (or NIE) for Greebles; both (a and b) were from Gauthier97 paradigm; (c) FFA NIE for faces; (d) for Greebles; (c and d) were from Gauthier98 paradigm; (e and f) the Summed-t values of NIE. T-values were summed and averaged across all FFA voxels corresponding to the upright vs. inverted stimulus version. (e) for Gauther97, (f) for Gauthier98. For all graphs, error bars denote standard errors.

Though the PSC is now more pervasively used in the fMRI papers, the summed-t values, which was originally reported in Gauthier et al. (1999), might be another possible index to verify NIE. This method sums the t-values from each participant’s FFA voxels, and then averages across upright minus inverted conditions (and averages across subjects as well). As each participant’s FFA contained different numbers of voxels, our summed-t values for faces and Greebles, again across 3 sessions, showed no significant interactions (though with trends of increase for Greebles NIE, in Gauthier97) (see Figure 9e and 9f).

#### Meta-analysis of NIE

One untested assumption in supporting the NIE as the index of the “face specificity of FFA” account is its reliability: to be a strong indicator of face-related processing, NIE has to be consistently identified, in relevant literature whenever upright vs. inverted faces were compared. To find out, we performed a search with the keywords: “FFA” and “neural inversion effect”, which yielded 18 fMRI articles matching the criteria of “fMRI studies containing FFA NIE results”. As shown in Table 1, seven articles identified positively significant NIEs, two articles found negatively significant NIEs (i.e. FFA activities for inverted faces higher than those for upright faces), while the remaining 9 articles showed no significant NIE for faces at the FFA.

A funnel plot of the 13 studies with reported stats was also shown in Fig. 10. Five of the aforementioned studies reporting non-significant results were excluded because they did not provide specific statistics (*p* or *t* value). Considering these exclusions, the meta-analysis results suggest that NIEs in the FFA suffered certain degrees of publication bias. While the meta-analysis (i.e. funnel plots) results are mostly related to the degree of publication biases, the current results do, however, suggest that the relatively skewed distribution of positively, null, and negatively significant NIE@FFA in the literature, seriously undermining its reliability assumption as face-specific index (i.e. most, if not all, published studies should yield consistently positively significant NIE@FFA for faces).

**Figure 10.**
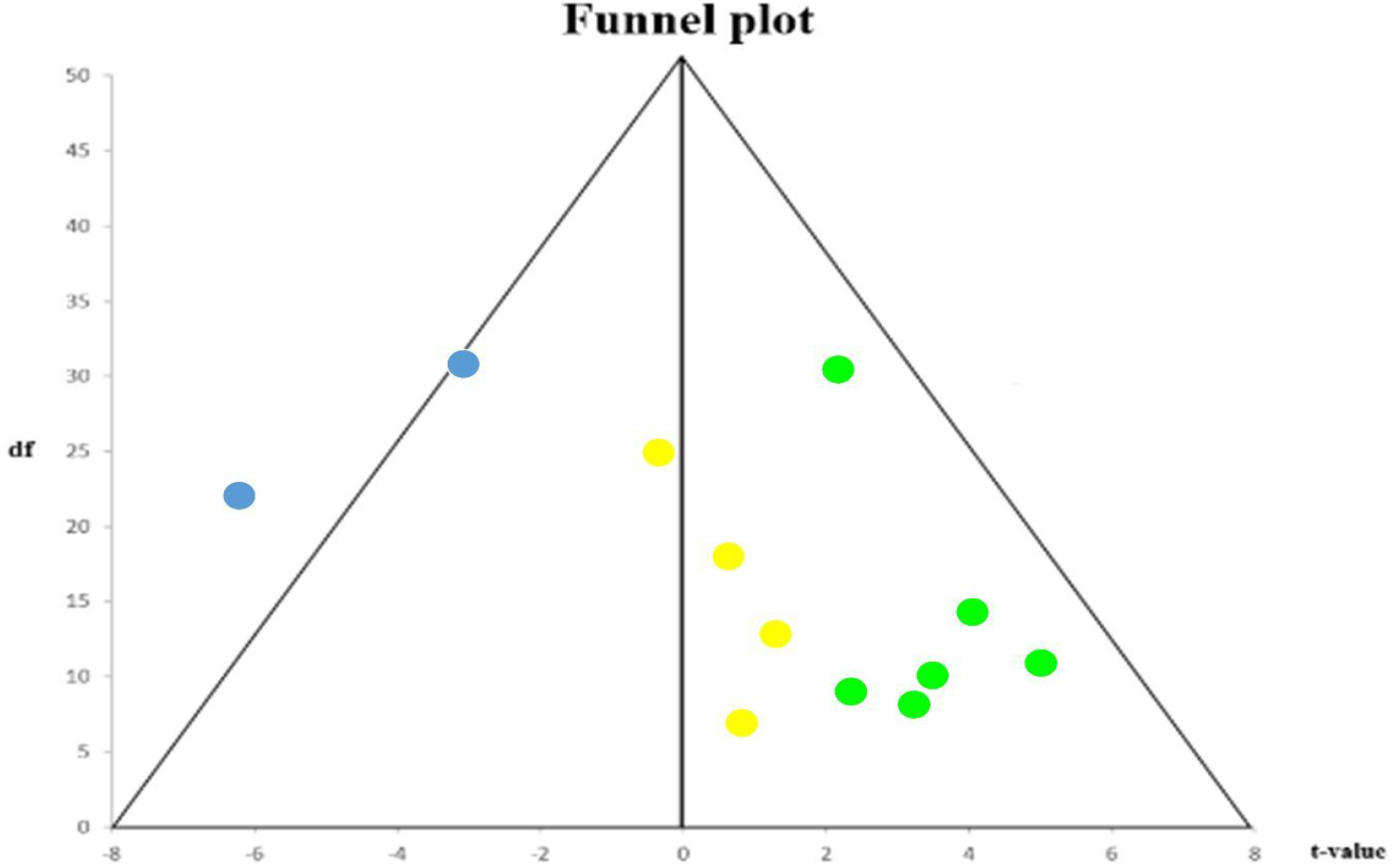
The funnel plot of reported statistics from the upper table. The unreported statistics in dark yellow (5 of 9), plus the 2 negatively significant NIE, all argue against the unidirectionality (i.e. only positively significant) of the neural inversion effect.

### (c) Neural Adaptation Effect in the FFA

Another index of successful training would be to see whether the beholders’ FFA responses for faces, after becoming Greeble experts, will become weaker (aka. adapted) following the presentation of Greebles, compared to the control condition of before-training session (where Greebles are not face-like to FFA, yet), and compared the inappropriate training condition (e.g. Gauthier97 vs. Gauthier98). Other than the two controlled comparisons, there is one more specific prediction: the neural adaptation would only happen to the faces in the “Faces after Greebles” condition (not the other way around, i.e. “Greebles after Faces”, because Greebles would always be adapted following faces in FFA, which is constantly face-selective), compared to the controlled “Faces after Objects” condition. Fig. 11 shows such t-test comparison results for both the before-vs. after-training sessions, and for both Gauthier97 and Gauthier98 regime.

**Figure 11.**
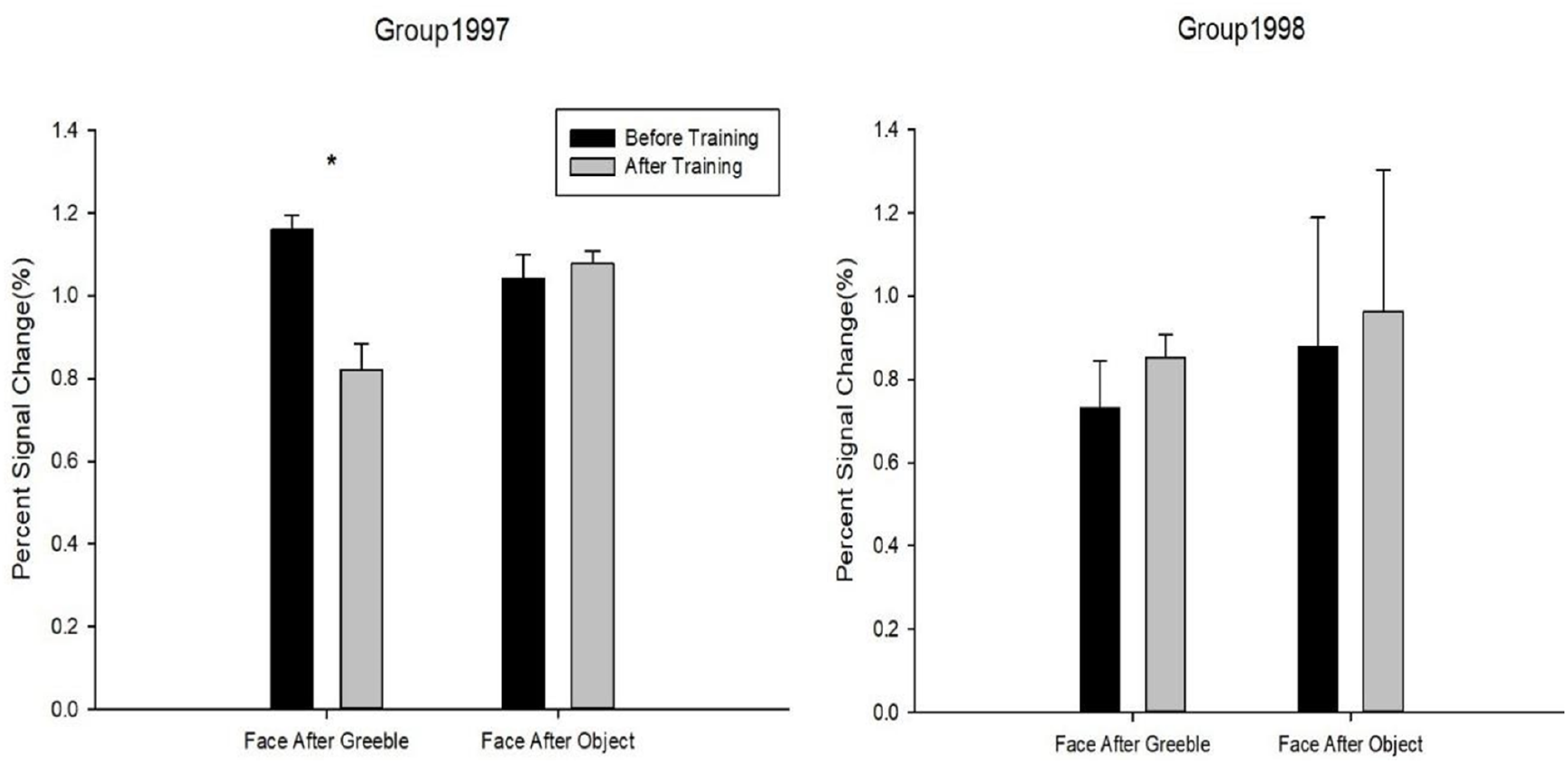
The neural adaptation effect in the FFA. The comparisons of percent signal changes for “Faces after Greebles” vs. “Faces after Objects” in both Gauthier97 (N=8) vs. Gauthier98 (N=8) training regime, were taken as the evidence for the support of perceptual expertise hypothesis of FFA. Error bars represent standard errors.

For the Gauthier97 group, there was a significant difference between the “Faces after Greebles” condition [*t*(7) = 2.01, *p* = 0.04], supporting the training-inducing Greeble expertise recruiting face-like responses in the FFA, thereby making the subsequent faces “adapted”. In contrast, the “Faces after Objects”, plus neither “Faces after Greebles” nor “Faces after Objects” comparisons in the Gauthier98 group, showed any adaptation effects in the FFA. Together, these 4 comparison results are in strong agreement with the predictions made by the perceptual expertise account of the FFA.

## DISCUSSION

The present work revisited two of the assumptions behind Brants et al. (2011) JOCN article, one of the seminal studies in the prolonged discussions for vs. against the perceptual expertise of FFA, the most prominent example in cognitive neuroscience. As one of the oft-cited stimuli in the field of cog-neuro, Greebles have been cited in many textbooks of vision research. And the claim that the training-induced FFA activity increases was primarily due to Greebles’ face-likeness, instead of the original putative expertise training, surely raises the concern about the premises of those assertion: that (1) Brants et al. (2011) also replicated the Greeble training effects, at least behaviorally; and (2) that the neural inversion effect, or NIE, was a reliable index of human face processing. As the current study has demonstrated, the first premise was unsubstantiated, that by differentiating the Greeble training paradigm as Gauthier97 vs. Gauthier98 version, the session-wise RT of the verification task between the two paradigm (Fig. 3) was drastically different, suggesting that Brants et al. (2011) adopted the Gauthier98 paradigm (by their almost identical RT results), whereas the original Greeble training study (i.e. Gauthier et al., 1999) adopted the Gauthier97 version. Further individual RT analysis in Gauthier98 (e.g. Fig. 5) showed that the large variability among participants was the main reason behind the in-convergibility of RTs for trained vs. untrained Greebles, the main difference between the original Gauthier98 training results and the current study.

In addition, the current study further provided three pieces of the fMRI evidence for the perceptual expertise of the FFA: (a) FFA activities to Greebles after training were significantly larger than that of the before-training condition, only under the appropriate (i.e. Gauthier97) training condition; (b) the lack of NIE effect in the FFA, the second premise for the Greebles-are-facelike hypothesis, plus the inconsistency of NIE effects at FFA in the extant literature (7 positive, 3 negative, and 8 no differences between upright vs. inverted faces) further nullified the backbone of the Brants et al. (2011) claim (that NIE@FFA is the reliable of human face processing); and (c) most importantly, the FFA adaptation effect, specifically for faces-after-Greebles, but not for faces-after-objects, and also not vice versa (i.e. Greebles-after-faces); under the appropriate (i.e. Gauthier97) training, and only after- (not before-, nor during-) training. Altogether, these combined evidences, including the clear RT difference by training paradigm and the close similarity between that of Gauthier98 (Fig. 3d) and Brants et al. (2011) at the behavioral side, and the three fMRI results (FFA training effect, no clear NIE@FFA, and FFA adaptation effect) on the neuronal side, the support for the perceptual expertise account, and against the face specificity account, of FFA were both presented.

The RT differences near the end of training in between the Gauthier97 and Gauthier98 paradigm were around 500 and 1000 ms, respectively. As depicted in Fig. 12, participants in the Gauthier98 training may have gone through more processing complications, which rendered the decision times of judging whether the image matched the label proportionally longer. The more the uncertainty during the decision processes (or more proposed steps), the longer (800 and 1200 ms for the named vs. unnamed Greeble versions in Gauthier98) the average reaction times at the end of training. These putative processes not only help explain the observed reaction time (RT) differences among different conditions, but also underlie the reason why Gauthier97 may be a relatively appropriate paradigm for fMRI effect of Greeble training.

**Figure 12.**
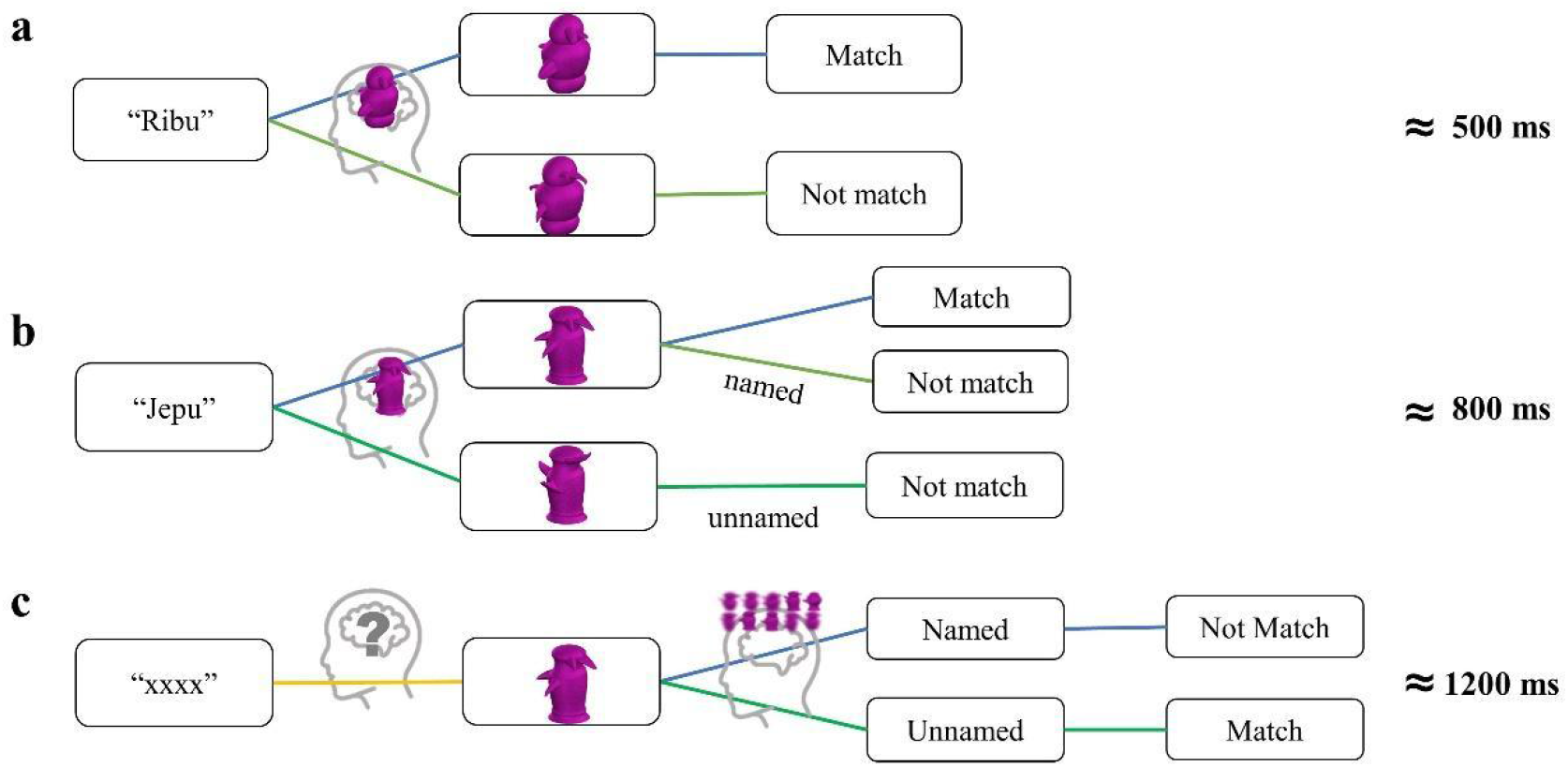
(a) The proposed task-flow of the verification task in the Gauthier97 training paradigm; (b&c) The same flowchart for the Gauthier98 training paradigm, for named vs. unnamed trials, respectively. The proposed steps in each condition (representing the putative stages of processing) proportionally correspond to the average response times at the end of training: Gauthier97 500 ms, Gauthier98 named trials: 800, and unnamed trials: 1200 ms. These differences may underlie the reasons why Gauthier97 is the relatively appropriate paradigm for Greeble training effects.

At the outset of this Greeble training study, the authors have struggled with the choice of the symmetry (e.g. symmetrical or asymmetric) of Greeble stimuli versions. Or final choice of using asymmetric Greebles was based on the following concern: the observed effects of FFA activities to Greebles, faces, or objects, if found as predicted, might still be attributed to the adage of “Greebles look like faces” (Farah, 2000), even though Greebles are the same facelike to both experts and novices (but FFA only respond significantly to the former after expertise acquisition). If a set of totally non-facelike stimuli, such as the symmetric Greebles of the present study, could be trained to drive FFA significantly after extensive training to enable automatic processing at the subordinate level, then the face-likeness may be less of an issue.

Despite this, it may still be argued that the current study has not replicated the results of Brants et al. (2011) and Gauthier et al. (1999), because of the difference of the Greeble versions. Our replies are: (a) our Gauthier98 behavioral results were similarly comparable to those in Brants et al. (2011), providing evidence that Brants et al. may have adopted this sub-optimal training paradigm, different from those in Gauthier et al. (1999) and the Gauthier97 of the current study. The similar behavioral response patterns (c.f. Fig. 3a and 3c to Fig. 3 of Gauthier97, as well as the Fig. 3b and 3d to Fig. 4 of Gauthier98, and Brants et al. Fig. 3), the dissimilarity among behavioral results from each paradigm, plus that the lack of literature support for the effect of stimulus asymmetry in short-term training (e.g., xxx), provide converging evidence against the existence of stimulus symmetry in affecting short-term training; (b) in terms of featural selectivity, FFA has been shown to prefer stimulus symmetry (Caldara et al., 2006), properties strongly associated with faces (Caldara & Seghier, 2009), but the effect (or trainability) of symmetry is yet to be quantified, thereby the effect of stimulus symmetry onto the training effect could be further assessed; and (c) currently there have been 4 studies comparing the training effects of both symmetric (Gauthier et al. 1999, and Brants et al., 2011) and asymmetric Greebles (Kung, Peissig, & Tarr, 2007, and the current study), while the effect of task (e.g., 1-back identity vs. passive viewing) have been shown to affect the observed training effect in the FFA (i.e. the 1-back task, due to its increased task demand, rendered the before-training FFA activities relatively higher, and relatively obscured the later increase of activities for training. Whether such cross-symmetry comparison could be made, could still be one of the potential research questions for future study.

Concerning future research questions, one of the possible extensions could be to incorporate the lasting effects of training into design. One general observation for Greeble experts, soon after their last after-training fMRI scan, was how soon they quickly forgot the associated image shapes and names (though there were also exceptions). Future training experiments, if adopted, could further investigate the extinction effects, and how soon they could be quickly recovered (and their associated brain mechanisms). Additional fMRI analysis methods, such as multivoxel pattern activities, representational similarities, or functional connectivities, or even deep neural network approaches, could all be viable options. Lastly, the effect of FFA adaptation, except its usefulness as a companion index of expertise acquisition, could also be tested in natural experts (such as bird or car experts), further extending its reliability and validity as an alternative, or even the primary, index of perceptual expertise.

## Acknowledgments

The authors would like to thank the NCKU MRI center for the support and consultation provided.

## Author Contributions

C.C.K. conceived the experiment. C.Y.C. developed the code, and K.L. carried out the experiment. K.L. and C.Y.C. analyzed the data. K.L. and L.S.W. prepared the figures. K.L., L.S.W. and C.C.K. discussed, revised, and completed the draft. All authors reviewed and approved the final manuscript.

